# A single neuron subset governs a single coactive neuron circuit in *Hydra vulgaris*, representing a prototypic feature of neural evolution

**DOI:** 10.1101/2020.11.22.392985

**Authors:** Yukihiko Noro, Hiroshi Shimizu, Katsuhiko Mineta, Takashi Gojobori

## Abstract

The last common ancestor of Bilateria and Cnidaria is believed to be one of the first animals to develop a nervous system over 500 million years ago. Many of the genes involved in the neural function of the advanced nervous system in Bilateria are well conserved in Cnidaria^1^. Thus, Cnidarian representative species, *Hydra*, is considered to be a living fossil and a good model organism for the study of the putative primitive nervous system in its last common ancestor. The diffuse nervous system of *Hydra* consists of several peptidergic neuron subsets. However, the specific functions of these subsets remain unclear. Using calcium imaging, here we show that the neuron subsets that express neuropeptide, Hym-176^2,3^ function as motor neurons to evoke longitudinal contraction. We found that all neurons in a subset defined by the Hym-176 gene (*Hym-176A*) or its paralogs (*Hym-176B*) expression^4^ are excited simultaneously, which is then followed by longitudinal contraction. This indicates not only that these neuron subsets are motor neurons but also that a single molecularly defined neuron subset forms a single coactive motor circuit. This is in contrast with the Bilaterian nervous system, where a single molecularly defined neuron subset harbors multiple coactive circuits, showing a mixture of neurons firing with different timings^5^. Furthermore, we found that the two motor circuits, one expressing *Hym-176B* in the body column and the other expressing *Hym-176A* in the foot, are coordinately regulated to exert region-specific contraction. Our results demonstrate that one neuron subset is likely to form a monofunctional circuit as a minimum functional unit to build a more complex behavior in *Hydra*. We propose that this simple feature (one subset, one circuit, one function) found in *Hydra* is a fundamental trait of the primitive nervous system.

## Main

The evolution of the nervous system in animals persists as one of the most intriguing and significant mysteries in modern biology. The first nervous system evolved in the last common ancestor of Cnidaria and Bilateria over 500 million years ago. Despite millions of years of divergence, an almost complete set of the neural genes needed to build the advanced nervous system found in bilaterians is well conserved in cnidarians^1^. This implies that the cnidarian nervous system is much simpler than the bilaterian nervous system, not because cnidarians have lost unnecessary neural functions during evolution, but rather because the ancient primitive nervous system in the common ancestor has long been sufficient for cnidarians to survive until the present day. Therefore, the cnidarian living fossil, *Hydra* is a good model organism for the study of the primitive nervous system in the last common ancestor and how it has evolved to the advanced nervous system.

The nervous system of *Hydra* is a net-like structure extending throughout the body and consists of multiple subsets of neurons, all of which are peptidergic. There is no clear evidence of other functional neurotransmitters being expressed in neurons. There are at least four neuropeptides, GLWa^6^, Hym-355^7^, RFa^8^ and Hym-176^2,3^, present in the *Hydra* nerve net (Fig.S1). The genes for GLWa and Hym-355 are co-expressed in the same neurons, which are scattered in the ectodermal layer all over the body. However, in the hypostome, there are some GLWa-positive and Hym-355-negative neurons together with the double-positive neurons^9,10^. RFa (*PreproA*)-*expressing* neurons are found in the tentacle, the hypostome, the upper body column and the peduncle (Fig.S1). The Hym-176 gene (*Hym-176A*) is expressed in the neurons of the hypostome, the body column and the peduncle. *PreproA* and *Hym-176A* are only co-expressed in the peduncle neurons^2,9^. Neither RFa-expressing nor Hym-176-expressing neurons are overlapped with GLWa/Hym-355 double-positive neurons^9,10^.

In addition to these observations, we recently demonstrated that the four-gene paralogs of *Hym-176A* are expressed in neurons in the tentacle (*Hym-176E*), the body column (*Hym-176B*), and the peduncle (*Hym-176C* and *Hym-176D*)^4^. *Hym-176B, Hym-176C*, and *Hym-176D* are all co-expressed with *Hym-176A*. Therefore, as summarized in Fig.S1, there are at least seven mutually exclusive molecularly defined neuron subsets (classical subsets) in the ectodermal layer in *Hydra* (subset I; GLWa/Hym-355, subset II; RFa (prepro A/C), subset III; RFa (prepro A/B), subset IV; Hym-176A/B, subset V; Hym-176A/C/D/RFa (prepro A), subset VI; Hym-176A/C/RFa (prepro A), subset VII; Hym-176E). Furthermore, a recent single-cell RNA-seq study of *Hydra*^11^ revealed a more precise molecular definition of nine neuron subsets (new subsets) in the ectodermal layer: subset I is further divided into subset ec3A, ec3B, and ec3C; subset III is further divided into subset ec4A and ec4B; both subsets V and VI form one single subset ec5; and the remaining subsets II/IV/VII correspond to subset ec2/ec1A/ec1B, respectively (Fig.S1).

Despite the comprehensive anatomical features of the *Hydra* nervous system, the function of the neuron subset is entirely unknown. The three neuron subsets (ec1A, ec1B and ec5) expressing *Hym-176A* and its paralogs (*Hym-176B, C, D, E*) cover the whole body of the animal in a regionspecific manner (ec1A in body, ec1B in the tentacle, and ec5 in the peduncle), suggesting that these neuron subsets are related to region-specific functions. Therefore, here we focus on these neuron subsets and examine their localized function by raising transgenic *Hydra* expressing the calcium indicator, GCaMP6s^12^, in each neuron subset.

### Functional characterization of neuron subsets expressing neuropeptide Hym-176 gene and its paralogs

We raised transgenic *Hydra* expressing the calcium indicator, GCaMP6s, in the neurons expressing each one of the *Hym-176* gene paralogs (*Hym-176A, B, C*, and *D*) under the control of the gene regulatory regions of these paralogs, as described previously^4^. The transgenic line, Hym-176B::GCaMP, visualized subset IV (ec1A) in the body column (Fig. S1). The transgenic line, Hym-176A::GCaMP, Hym-176C::GCaMP and Hym-176D::GCaMP, all visualized the same subset, i.e., subset V and VI (ec5) in the peduncle. We were unable to visualize subset VII (ec1B) in the tentacle due to the difficulty in generating the transgenic line, Hym-176E::GCaMP.

These subset-specific GCaMP6s expressions showed clear excitation patterns in each neuron subset (Movie S1-S4). The shape of the blinking neurons and their resulting net-like structure was especially visible in the close-up view (Movie S5). We quantified the timing and the intensity of the excitation visible in the recorded movies. For example, we selected 18 neurons from the *Hym-176A*-expressing subset (ec5) in Movie 1 (Fig. 1A); their excitation patterns for 15 sec are shown (Fig.1B and S2A). Each of the 18 neurons fired simultaneously at 2.22, 4.79, 6.81, 9.38, and 12.50 sec (vertical dotted lines). The simultaneous firings in this subset (ec5) and the other subset (ec1A) were also visualized by the transgenic lines, Hym-176C::GCaMP (Fig. 1E/1F/S2C, Movie 3/S3), Hym-176D::GCaMP (Fig. 1G/1H/S2D, Movie 4/S4) for ec5, and Hym-176B::GCaMP for ec1A (Fig. 1C/1D/S2B, Movie 2/S2). This simultaneous firing was confirmed by the distribution of the spike timing between all the tested neurons (Fig. 2B/E/H/I). For more than 80% of spikes (22/27), the spike timing of the tested neurons in the IQR (interquartile range) varied by less than 0.06 sec. The neurons in the bud were all excited at the same time but different from those in the parental polyp (cell# 41,42, 45, 46, 49, and 54 in Fig. 1F/S2C). These results suggest that each of the two neuron subsets (ec1A and ec5) forms a single coactive circuit and that the circuit in the bud at this developmental stage is independently regulated from the circuit in the parental polyp.

**Fig. 1.**
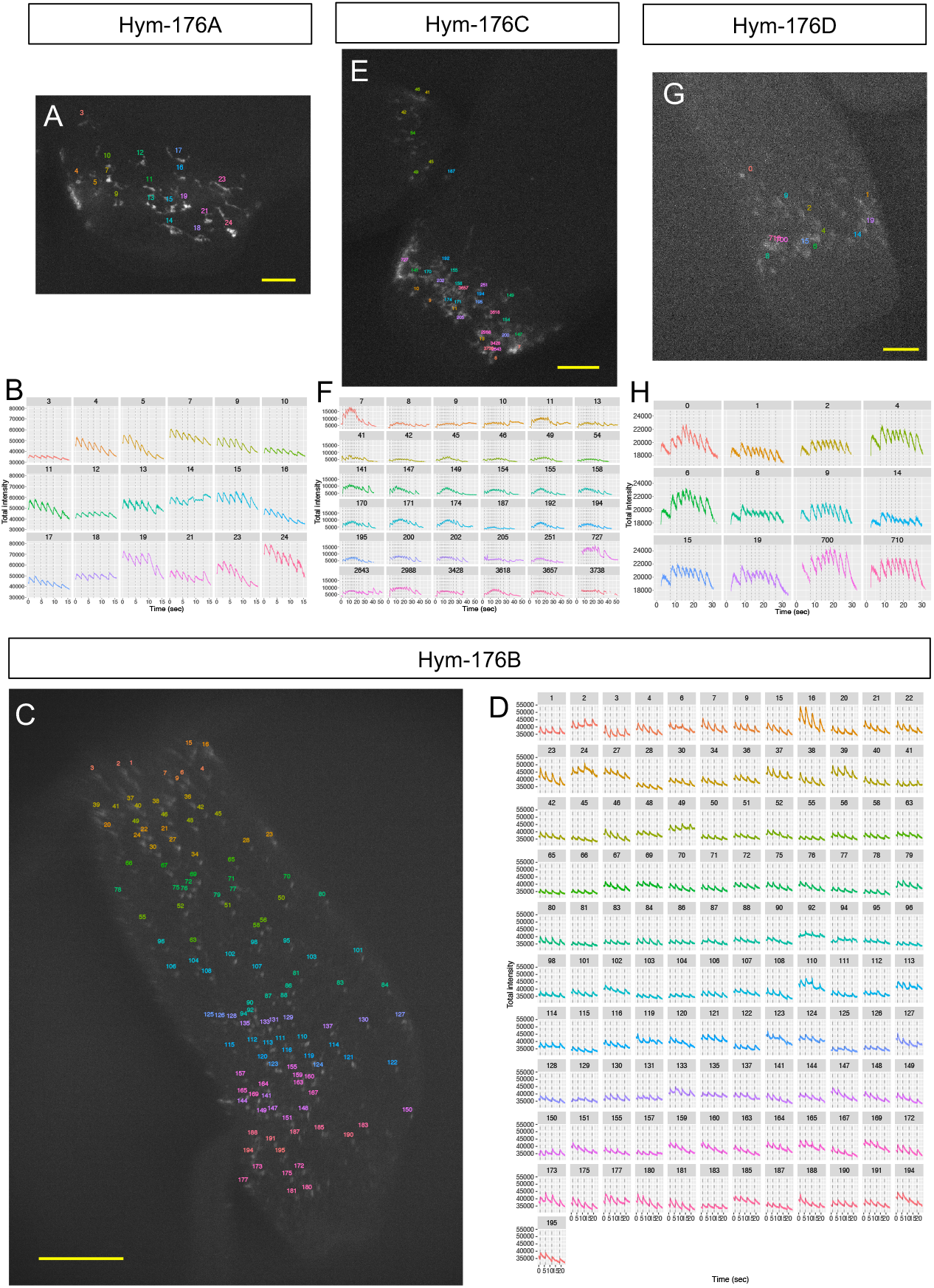
Simultaneous neuronal excitation of the peptidergic neuron subsets expressing neuropeptide gene *Hym-176* paralogs. *Hym-176A-expressing* neuron subset (A, B). *Hym-176B-* expressing neuron subset (C, D). *Hym-176C-expressing* neuron subset (E, F). *Hym-176D-expressing* neuron subset (G, H). Position of tested neurons (A, C, E, G). Neuronal activity (total intensity of GCaMP) is shown, with the vertical dashed lines indicating the average starting time of excitation of all tested neurons in each subset (B: 2.22, 4.79, 6.81, 9.38, 12.50 sec; D: 1.09, 5.75, 11.72, 18.59 sec; F: 2.13, 5.57, 7.98, 9.72, 11.60, 13.00, 14.83, 16.94, 19.45, 22.27, 26.26, 33.86 sec; H: 7.37, 10.70, 13.37, 15.32, 17.41, 19.77, 22.80, 26.15 sec) except neuron #41-54 in F (see text). The number in the strip and the color of the excitation profile for each subset correspond to cells in each one of A, C, E and G. Bar: 30 μm in A, 200 μm in C, 100 μm in E and 50 μm in G.

**Fig. 2.**
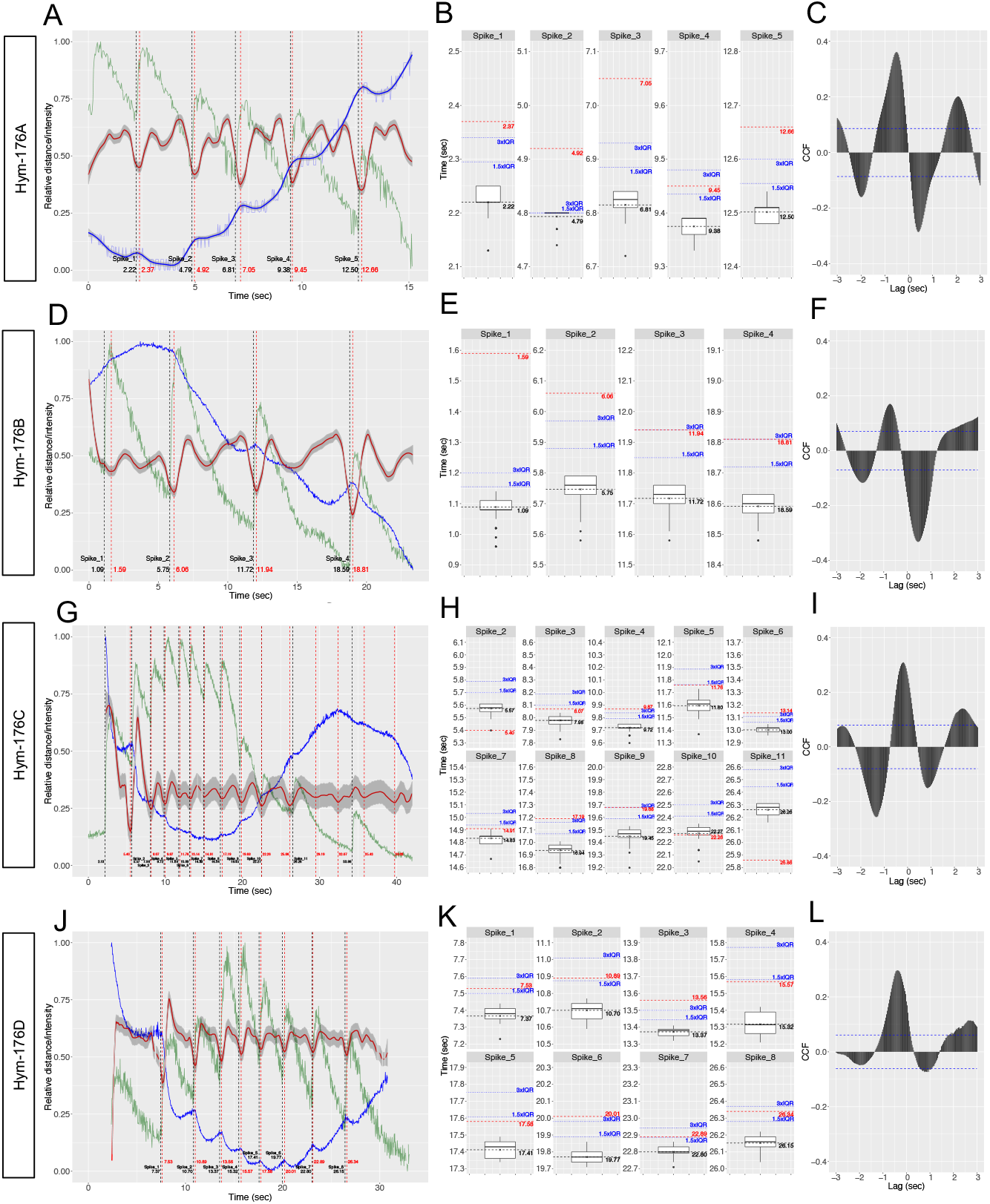
Neuronal activity in the Hym-176 peptidergic neuron subsets is associated with longitudinal contraction. *Hym-176A*-expressing neuron subsets (A, B, C). *Hym-176B*-expressing neuron subsets (D, E, F). *Hym-176C*-expressing neuron subsets (G, H, I). *Hym-176D*-expressing neuron subsets (J, K, L). Representative neuronal excitation in green; cell #24 in Fig. 1B (A), cell #16 in Fig. 1D (D), cell #7 in Fig. 1F (G) and cell #0 in Fig. 1H (J). Vertical dashed black lines show the spike train (A, D, G, and J), as shown in Fig. 1B, D, F, and H, respectively. Change in the distance between the two designated neurons in each subset is shown in blue (A, D, G, J); cell #14 and #17 in A, cell #3 and #150 in D, cell #13 and #149 in G, cell #0 and #8 in J. Second-order differences of the distance in red with 95% confidence interval (gray). Starting points of contraction (vertical red dashed line; A: 2.37, 4.92, 7.05, 9.45, 12.66 sec; D: 1.59, 6.06, 11.94, 18.81 sec; G: 5.40, 8.07, 9.87, 11.76, 13.14, 14.91, 17.19, 19.68, 22.26, 25.86 sec; J: 7.53, 10.89, 13.56, 15.57, 17.58, 20.01, 22.89, 26.34 sec). Box plot of the spike timing of all tested neurons for each spike (B, E, H, and K). The horizontal dashed black line indicates average spike timing, corresponding to the vertical dashed black line in A, D, G, and J. The horizontal dashed red line indicates the start of contraction, corresponding to the vertical dashed red line in A, D, G, and J. The horizontal dotted blue line indicates Q3 (3-quantile) + 1.5/3.0 x IQR (interquartile range). Cross-correlation (C, F, I, and L) between representative neuronal excitation (solid green line) and longitudinal contraction (solid red line) in A, D, G, and J, respectively. The horizontal dashed blue line indicates the 95% confidence interval cut-off.

Although the timing of the firing was the same between neurons in a subset, the intensity of the firing and its time course profile varied between the neurons, *e.g*., high (cell #19, #24) or low intensity (cell #3, #10), and steady (cell #12, #18) or decaying (cell #4, #5, #23, #24) or atypical (cell #14) oscillation in the *Hym-176A*-expressing neuron subset, ec5 (Fig. 1B/S2A). We observed these differential excitation profiles in the same subset with different transgenic lines (Fig. 1F/1H/S2C/S2D) and also in the different subset, ec1A (Fig. 1D/S2B). These results indicate that neurons in a coactive circuit respond synchronously but differentially.

Quantitative analysis also demonstrated that the simultaneous excitation of neurons in each of the two subsets, ec1A and ec5, was well correlated with or was mostly followed by the longitudinal contraction. We evaluated this correlation by measuring the distance between the two designated neurons, usually the uppermost and the lowermost neuron in each subset (blue lines in Fig. 2A/D/G/J). Second-order differences in the distance (red lines in Fig. 2A/D/G/J) indicate a contraction status, i.e., fully relaxed at the local minimum and completely shrunk at the local maximum. Thus, fully relaxed at the local minimum indicates the start of contraction (vertical dashed red line in Fig. 2A/D/G/J). Cross-correlation between the representative neuron excitation (green line in Fig. 2A/D/G/J) and the second-order difference of the distance was above the 95% confidence interval cut-off (horizontal dashed blue line in Fig. 2C/F/I/L). Most (24 out of 27) of the neural excitations (vertical dashed black lines) were followed by longitudinal contractions (vertical dashed red lines) in less than 0.4 sec (Fig. 2A/D/G/J). This delay between spike and contraction was far beyond the dispersion of spike timing among all tested neurons (median IQR: 0.06 sec). For instance, for more than 74% (20/27) of spikes, contraction started later than Q3 (3-quantile) + 1.5xIQR. For more than 51% (14/27) of spikes, contraction started later than Q3 + 3xIQR (Fig. 2B/E/H/K). These results suggest that the dispersion of spike timing is minimal (almost at the same time) among the tested neurons in a subset, compared to the timing of the start of the contraction, and that contraction indeed followed the preceding simultaneous excitation. These results demonstrate that the neuron subsets, ec1A and ec5 expressing the *Hym-176* gene paralogs, function as coactive motor neuron circuits that evoke longitudinal contraction.

Besides those contractions associated with the preceding simultaneous excitation, we found some of the contractions were not associated with excitation (*e.g*., the contractions at 29.19, 32.07, 35.43 and 39.33 sec in Fig. 2G). We do not yet entirely understand these unassociated contractions, but they may result from responses to the residual neurotransmitter released by the preceding contraction-associated excitation. The contraction-unassociated excitation at 33.86 sec in Fig. 2G may not have been able to evoke contraction due to epitheliomuscular desensitization of the neuronal excitation.

### Two independent motor neuron circuits coordinately function to regulate their region-specific contractions

Since both of the two coactive motor neuron circuits, ec1A in the body column and ec5 in the peduncle, evoked longitudinal contraction, we examined whether these two circuits form a single coactive circuit or they function independently. At the onset of the continuous longitudinal contraction in a double transgenic line or an operative chimera of single transgenic lines, we found that all the tested neurons in the peduncle (foot) neuron circuit only were simultaneously excited at their first spike (cell #13-#19 at 0 sec in Fig. 3A/3B/S3A, Movie 5, cell #19-#89 at 6.42 sec in Fig. 3C/3D/S3B and Movie 6). In contrast, those in the body neuron circuit were mostly silent, except for only a few neurons (cell #1 and #6 in Fig. 3D). This suggests that the two circuits were independently regulated without forming a single coactive circuit. Furthermore, with the spike only found in the foot circuit, 2-4 times stronger contraction of the foot than the body column was evoked (green arrows, at 0 sec in Fig. 4A and 6.5 sec in Fig. 4C). This indicated that the foot motor circuit was excited independently of the body motor circuit and evoked mostly foot contraction, at least at the onset of the continuous longitudinal contraction. Subsequent contractions were synchronized again between the two circuits, following their synchronized excitation (Fig. 4C). However, sometimes subsequent contractions were anti-synchronized, especially when the foot was attached to the substrate, where it seemed like the foot tried to contract in vain because the body contraction simultaneously hampered it (Fig. 4A). These repelling contractions also indicate the independent motor function of these two circuits.

**Fig. 3.**
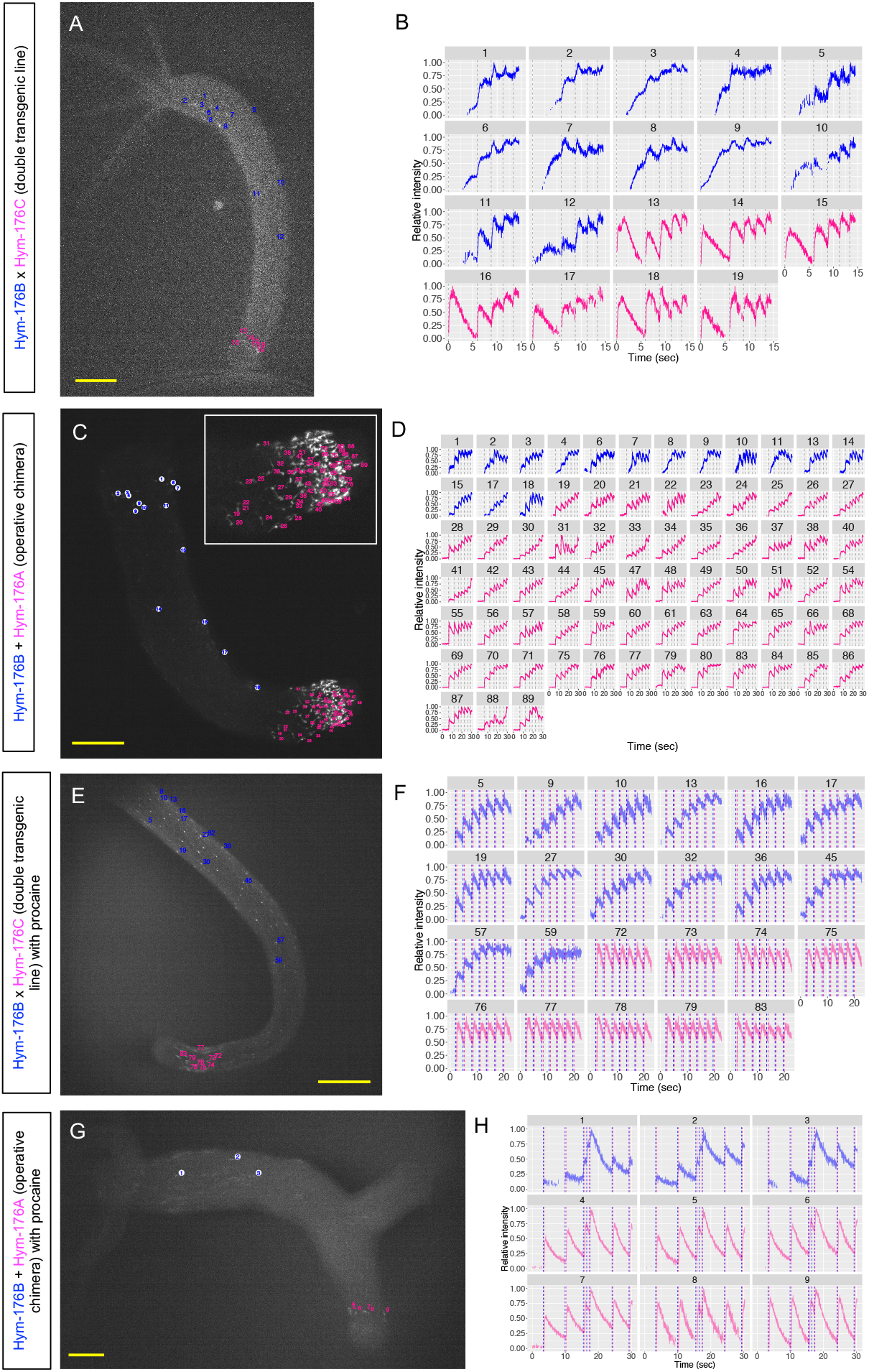
Simultaneously visualized neuronal activity of body and foot circuit. Double transgenic line of Hym-176B::GCaMP and Hym-176C::GCaMP with (E, F) or without (A, B) procaine treatment. The operative chimera of Hym-176B::GCaMP and Hym-176A::GCaMP with (G, H) or without (C, D) procaine treatment. Position of tested neurons (A, C, E, G). Neuronal activity (normalized intensity of GCaMP) with vertical dashed black lines indicating coactive neuronal excitation of both circuits (B: 0, 5.84, 8.71, 11.11, 13.19 sec; D: 6.42, 11.99, 16.28, 19.08, 22.31, 25.61, 28.85 sec). Neuronal activity (normalized intensity of GCaMP) with vertical dashed lines indicating coactive body neuron excitation in blue (F: 1.94, 4.74, 7.94, 10.66, 13.32, 16.40, 19.42 sec; H: 3.25, 9.84, 15.30, 16.31, 17.28, 23.92, 29.00 sec) and coactive foot neuron excitation in red (F: 2.39, 5.16, 8.42, 11.11, 13.79, 16.78, 19.81 sec; H: 3.53, 10.09, 15.65, 17.16, 24.13, 29.21 sec). The number in the strip and the color of the excitation profile for each animal correspond to cells in each one of A, C, E and G (body neuron circuit in blue and foot neuron circuit in red). Bar: 100 μm in A, E, G and 200 μm in C.

**Fig. 4.**
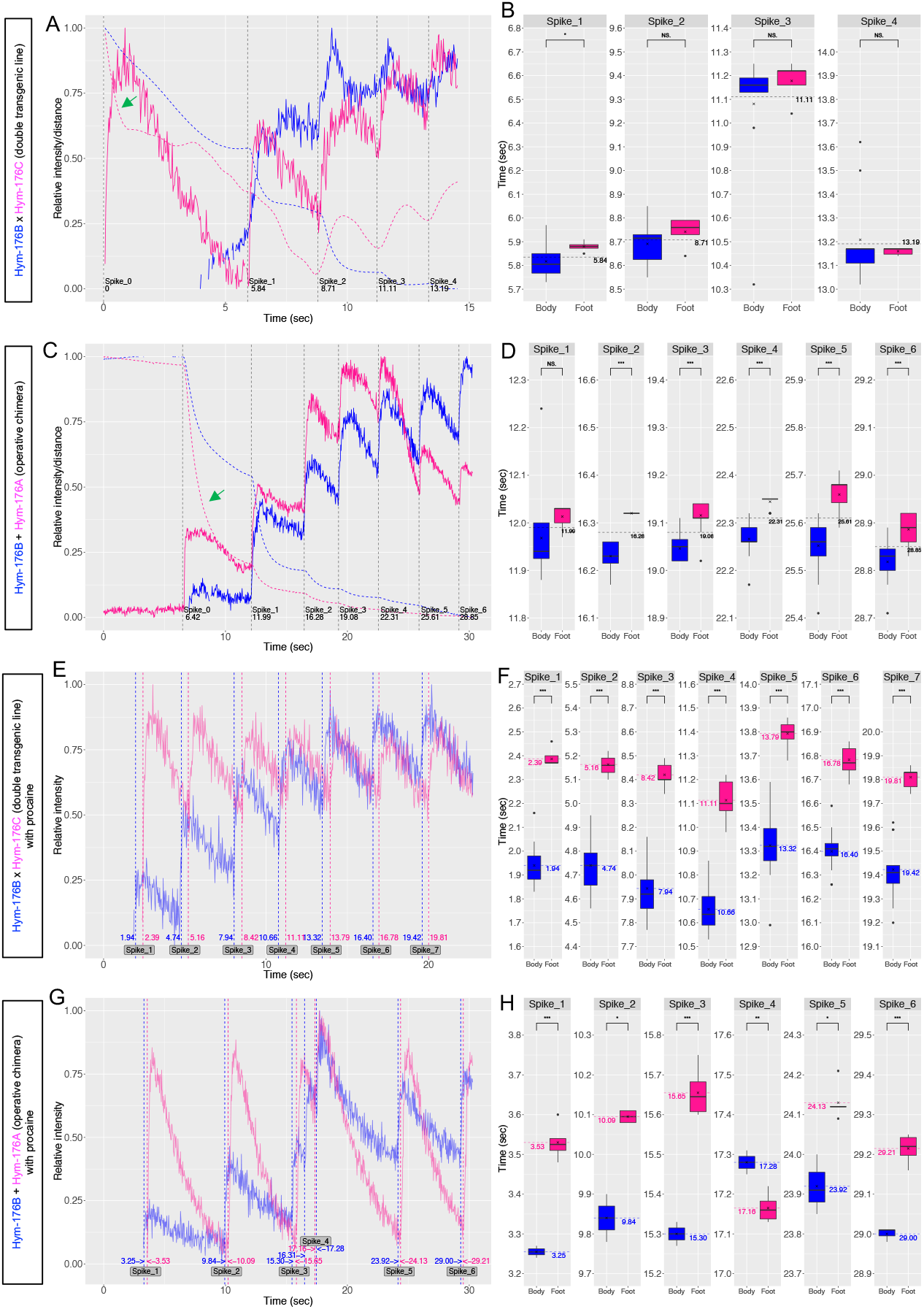
Coordinated interaction between body and foot neuronal circuit. Double transgenic line of Hym-176B::GCaMP and Hym-176C::GCaMP with (E, F) or without (A, B) procaine treatment. The operative chimera of Hym-176B::GCaMP and Hym-176A::GCaMP with (G, H) or without (C, D) procaine treatment. Normalized neuronal activity of representative neurons from body circuit in blue and foot circuit in red; cell #1 and #16 in Fig. 3B (A), cell #15 and #89 in Fig. 3D (C), cell #5 and #83 in Fig. 3F (E), cell #2 and #9 in Fig. 3H (G). Vertical dashed black/blue/red lines are as described in Fig. 3. Normalized distance between the three designated neurons in each animal; cell #9, #14, and #16 in Fig. 3B (A), cell #7, #21, and #87 in Fig. 3D (C). Distance between the first two neurons, shown by the dashed blue line, and between the last two neurons, shown by the dashed red line, indicates body and foot length, respectively. Box plot of the spike timing of all tested neurons in the body (blue) and foot (red) circuit for each spike (B, D, F, H). The horizontal dashed black line in B and D indicates the average spike timing of all tested neurons in both circuits, corresponding to the vertical dashed black line in A and C, respectively. The horizontal dashed blue/red line in F and H indicates the average spike timing of all tested neurons in the body/foot circuit, corresponding to the vertical dashed blue/red line in E and G, respectively. Statistical significance of the difference between average spike timings of body and foot circuit was evaluated by two-sided Student’s t-test; NS: no significant difference; *: p < 0.05; **: p < 0.01; ***: p < 0.001.

Although only the foot circuit was excited at the onset of the continuous contraction, the subsequent excitations were completely synchronized between the body and the foot circuits. We found that procaine, a voltage-gated sodium channel blocker, uncoupled the synchronized firing of these two circuits. The double transgenic line or the operative chimera of single transgenic lines treated with 1% procaine stopped contraction but maintained excitation. We observed a phase shift between the two coactive circuits: body one (blue) first and foot one (red) next (Fig. 3E-H, 4E-H, S3C/D and Movie 7, 8), although sometimes foot one at 17.16 sec preceded the body one at 17.28 sec (spike_4 in Fig. 3G/H). These inter-circuit phase shifts (0.435 sec on average for Fig. 4E/F and 0.262 sec on average for Fig. 4G/H) were significantly larger than the intra-circuit dispersion of spike timing (0.087 sec and 0.038 sec on average, respectively). These shifts were not observed in the absence of procaine (Fig. 4B/D). We also found an irregular spike in the body neuron circuit at 16.31 sec (Fig. 4G). All these results suggest that the two coactive motor circuits are not in a single coactive circuit but are coordinately regulated, resulting in the inter-circuit synchronicity.

### Light-activated signaling center and unidirectional neurotransmission in the body neuron circuit

Besides the inter-circuit phase shift described above, we found that the prolonged procaine treatment also shifted the intra-circuit phase of excitation. For each of the four spikes, when we plotted the spike timing of all the tested neurons in the body circuit against the position of the neurons along the body axis, numbered in order beginning from head to foot, the cells closer to the lower end of the body column (in pink) were then excited later than the cells below the head (in blue) (Fig. 5A-C). This suggested that the wave of excitation transmitted unidirectionally from just below the head down to the foot (Movie 9). This unidirectional neurotransmission was probably too quick to be detected without the procaine, which slowed down the transmission.

**Fig. 5.**
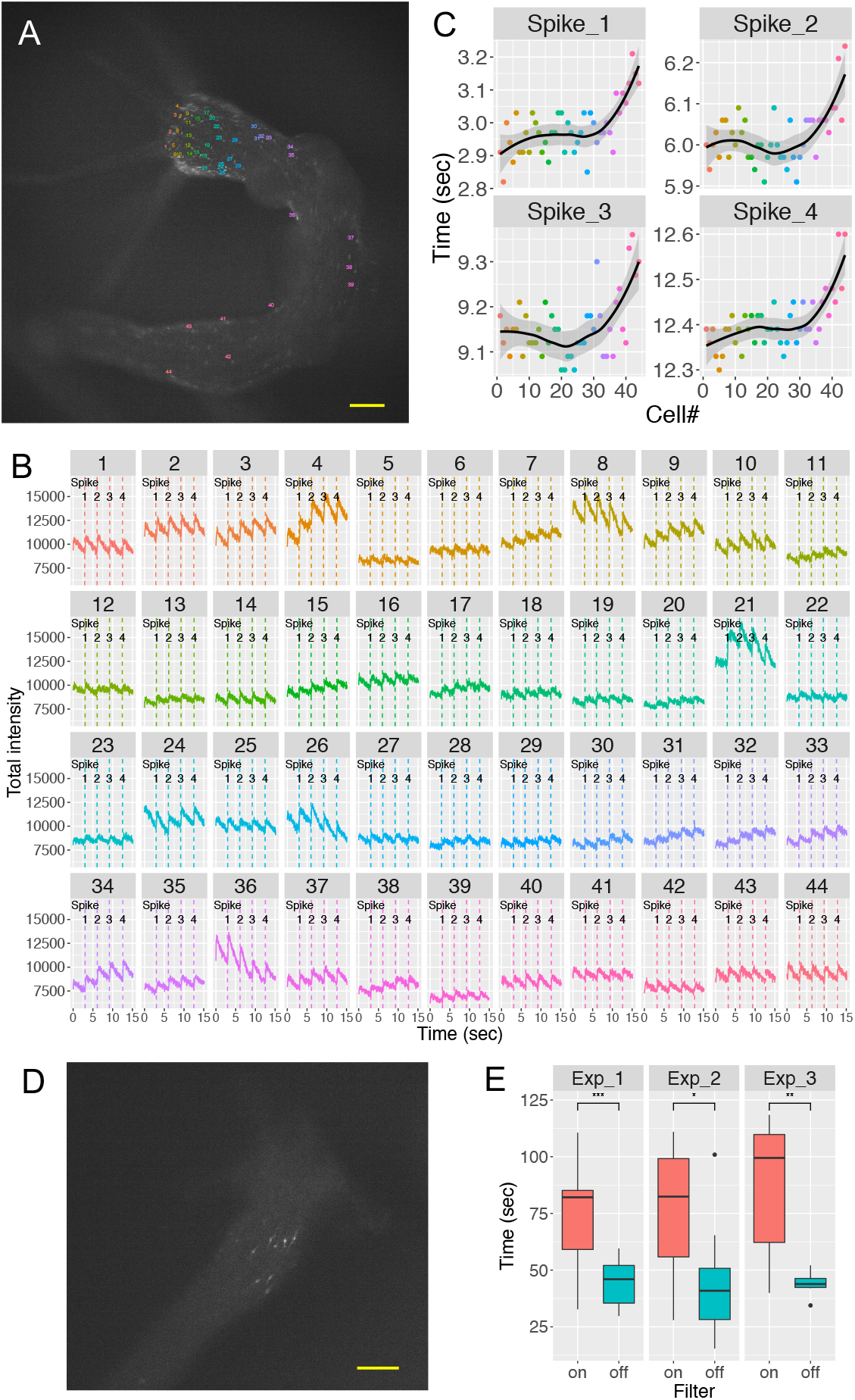
Unidirectional signal transmission in the body neuron circuit. Hym-176B::GCaMP transgenic animal was treated with procaine. Tested neurons are numbered according to their position, ascending from head to foot (cell#) (A). Neuronal activity (total intensity of GCaMP) with vertical lines indicating individual excitation spikes of each neuron (B). For each spike, the starting time is plotted against the cell# (C). The number in each strip, the color of the excitation profile in B, and the color of dots in C correspond to cells in A. The solid black line indicates loess regression with the 95% confidence interval in gray. Light-activated neurons in Hym-176B::GCaMP transgenic animal (D). Bar: 100 μm both in A and D. Box plot of time, which was taken by light-activated neurons to start excitation when exposed to blue light with or without dark filter (E). Three independent experiments were conducted (n = 22 for exp_1 and 2, and n = 14 for exp_3). Statistical significance of the difference between the average time to the beginning of excitation with and without a dark filter was evaluated using two-sided Student’s t-test; NS: no significant difference; *: p < 0.05; **: p < 0.01; ***: p < 0.001.

Although the body neuron circuit is a single coactive motor circuit, as we mentioned above, we found that a group of neurons in the tentacle base and around the neck were excited earlier than the rest of the neurons in the circuit at the onset of the body contraction (Fig. 5D, Movie 10). This preceding excitation occurred more easily with more intense (off-filtered) blue light for the excitation of GCaMP (Fig. 5E), suggesting these neurons responded to light as a part of a light-sensing system to trigger the excitation of the motor neuron circuit and subsequent longitudinal contraction. We believe these neurons belong to a subset different to the subset ec1A because detecting light or mediating the light-sensing requires an additional gene expression.

Taken together, the light-activated neurons near the head function as a signaling center to integrate light stimulation, and the body neuron circuit transfers the signal unidirectionally down to the foot.

## Discussion

Before the calcium imaging technique became widely available, functional analysis of the nervous system in *Hydra* was quite limited, except for a series of pioneering studies by Josephson and Passano^13,14^, who used intracellular electrical recordings. They showed that there are two types of action potentials in *Hydra:* one type is correlated with movement (*e.g*., contraction burst (CB) and tentacle pulse) while the other type is not correlated with movement (*e.g*., rhythmic potential (RP)). In their first successful attempt at calcium imaging of the *Hydra* nervous system, Dupre and Yuste functionally identified almost all of the neurons in *Hydra*. They categorized them into five coactive circuits, some of which seemed to correspond to the aforementioned action potentials. They named the five coactive circuits CB, STN, RP1, RP2, and Others^15^. However, the molecular identities of these circuits remain unknown.

In this study, we characterized subset-specific neuronal activity to reveal the relationship between functionally identified circuits and molecularly identified subsets. These two are not always identical. We found that each one of the two peptidergic neuron subsets (one expressing *Hym-176B* in the body column (ec1A) and the other expressing *Hym-176A/C* in the peduncle (ec5)) is a coactive motor neuron circuit because the simultaneous excitations of almost all neurons in each subset were followed by longitudinal contractions (Fig. 1, 2, Movie 1-4). This finding demonstrates that a subset consists of a single circuit in *Hydra* (Fig.6). In contrast, multiple circuits usually share a molecularly defined neuron subset in the advanced nervous system^5,16^. Therefore, we propose here that this notion of “one subset, one circuit” is a characteristic feature of the primitive nervous system.

**Fig. 6.**
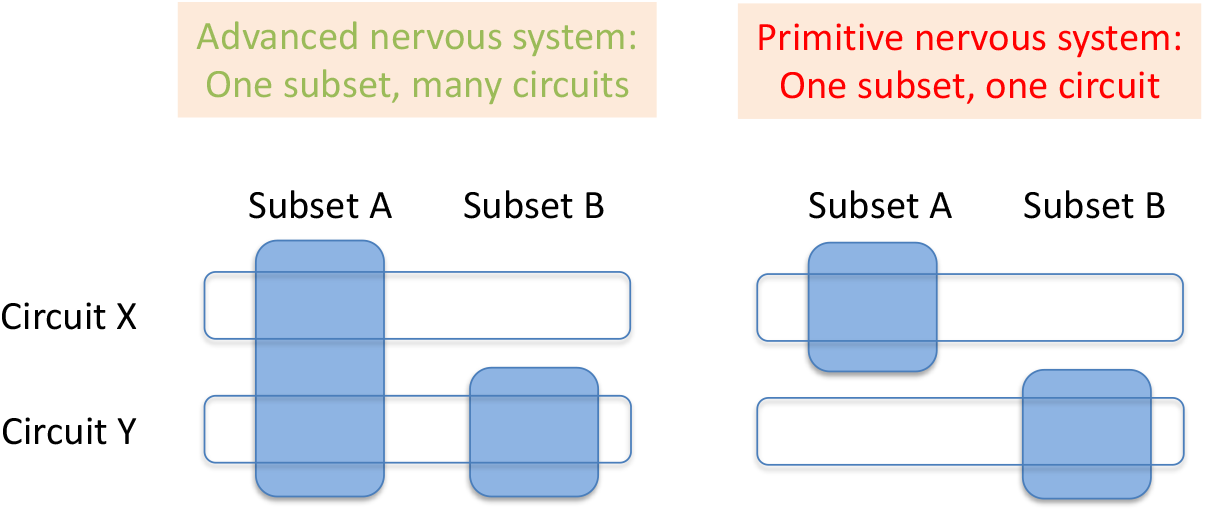
One subset, one circuit in the primitive nervous system. Neuron subset is a molecularly defined group of neurons expressing a common marker gene set. A neuron circuit is a physiologically defined group of neurons, firing simultaneously to function. Usually, a neuron subset harbors multiple neuron circuits in an advanced nervous system while a neuron subset harbors a single neuron circuit for their motor function in *Hydra*.

The circuit CB is the only circuit that Dupre and Yuste identified to be correlated with the longitudinal contraction. The circuit is distributed throughout the whole body except in the tentacle and the basal disk^15^. Their findings on the circuit CB appear to correspond to the two motor circuits that we identified in this study, even though the CB is one single coactive circuit. This may be because the two motor circuits are closely associated temporally. We found that procaine is useful to distinguish the two motor circuits, demonstrating that they are indeed two circuits that are synchronously regulated (Fig. 3, 4). This inter-circuit synchronicity was broken at the onset of continuous contraction to exert region-specific motor function (Fig. 4A, C). Alternatively, even with synchronous firing, subsequent contractions were not always synchronized between the circuits due to a physical restriction that prevented simultaneous contraction of the body and the peduncle (Fig. 4A). These findings demonstrate that the two circuits function independently as region-specific motor neuron circuits. Therefore, we expand the notion mentioned above to the following upgraded notion: “one subset, one circuit, <u>one function</u>”.

The coordinated regulation of the region-specific motor neuron circuits may be required to build a more complex behavior, such as somersaulting in *Hydra*^17^, in which several simple behavioral units are sequentially executed. Standing animals first attach to the floor with their tentacles (step 1), detach the foot from the floor (step 2), contract the body column to find a new place (step 3), and finally attach the foot to a new place (step 4). The initial foot-specific contraction we observed at the onset of the continuous longitudinal contraction may reflect the second and third steps of somersaulting (Fig. 3A-D, 4A-D). Responding to light with the circuit in the head and the unidirectional neurotransmission from head to foot (Fig.5) implies how light may be one of the triggers for this complex movement.

In summary, we demonstrated that a molecularly defined peptidergic neuron subset serves as a single coactive motor neuron circuit in the manner of “one subset, one circuit, one function”. This would have been a fundamental trait in primitive nervous systems and represents an underlying mechanism to construct more complex behaviors by assembling functional subsets as behavioral units.

## Acknowledgment

This work was supported through funding from King Abdullah University of Science and Technology (KAUST) under award numbers BAS/1/1059/01/01 and URF/1/1976/03/01.

## Competing interests

The authors declare no competing interests.

## Data availability

Requests for materials should be addressed to Y.N or T.G.

## Author contributions

Y.N., H.S., and T.G. designed the research. Y.N. performed most of the experiments. H.S. raised and maintained transgenic lines together with Y.N.. K.M. contributed a lot to the computational works done by Y.N.. All the authors wrote the manuscript.

## Materials and methods

### Animals and culture

The transgenic strain AEP of *Hydra vulgaris* was used in all experiments, which were raised and cultured as described below or previously ^**1,2**^.

### Transgenic reporter lines

Transgenic *Hydra* was generated using a modified method derived from Wittlieb et al.^3^ as described previously^1^. Briefly, we replaced GFP in the previously reported expression vectors (Hym-176A::GFP; Acc# LC426372, Hym-176B::GFP; Acc# LC426373, Hym-176C::GFP; Acc# LC426374, Hym-176D::GFP; Acc# LC426375) with the calcium indicator, GCaMP6s (#40753, Addgene, UK), so that it could be expressed in the neuron subsets expressing each one of the *Hym-176* gene paralogs (*Hym-176A, B, C*, and *D*). The 1- to 2-cell stage of embryos were microinjected with the vectors. The hatched F0 polyps were screened to select the GCaMP positive polyps, which were then mated to each other or wild AEP to obtain F1 transgenic lines. The transgenic lines, Hym-176B::GCaMP and Hym-176C::GCaMP were F1. The transgenic lines, Hym-176A::GCaMP and Hym-176D::GCaMP were F0. The double transgenic line (Hym-176B::GCaMP x Hym-176C::GCaMP) were obtained by crossing F1 polyps of Hym-176B::GCaMP and Hym-176C::GCaMP. The operative chimera of Hym-176B::GCaMP and Hym-176A::GCaMP (Hym-176B::GCaMP + Hym-176A::GCaMP) was made by grafting the upper half of the Hym-176B::GCaMP line and the lower half of the Hym-176A::GCaMP line.

### Signal detection and quantitative analysis of excitation and longitudinal contraction

Transgenic animals were sandwiched between two slide glasses with a spacer (0.1 mm thick). The GCaMP signals were detected under the fluorescent dissection microscope (SMZ25, Nikon, Japan) equipped with epi-fluorescence attachments and a CMOS camera (ORCA-Flash4.0 v2, Hamahoto, Japan). Neuron firings visualized by GCaMP were recorded at the speed of 30 msec/frame and tracked frame by frame using the Fiji^4^ plugin, TrackMate^5^ with its default setting (LoG detector; msb: 15-50, threshold: 0.01-4, and Simple LAP tracker). We manually edited the automated tracking results afterwards. The tracking data (intensity and position of each neuron at each frame) were analyzed by R^6^ and visualized by its graphic package, ggplot2^7^. The relative intensity was calculated by normalizing the total intensity with min-max feature scaling. The longitudinal contraction was evaluated as the change in the distance between two designated neurons through all frames, usually the uppermost and the lowermost neuron in each neuron subset. The distance was also normalized by min-max feature scaling. The second-order difference of the normalized distance was used to determine the start of contraction at its local minimum. Cross-correlation was calculated with function ccf in R. To compare the firing timing between body and foot circuits (Fig. 4B/D/F/H) or with and without attenuation of blue light (Fig. 5E), we carried out two-sided Student’s t-test using the R package, ggsignif. Boxplot elements are defined as follows: center line, median; cross mark, mean; box limits, the first and third quartiles (the 25th and 75th percentiles); whiskers, 1.5x interquartile range; points, outliers (data beyond the end of the whiskers).

### Procaine treatment

Animals were soaked in 1% procaine hydrochloride (P9879, Sigma-Aldrich) in a culture medium. The voluntary longitudinal contraction was stopped after a few minutes while the voluntary excitation was intact. The inter-circuit phase shift of excitation was first observed. Then, the intra-circuit phase shift was observed with prolonged incubation. This effect of procaine was reversible because animals contracted once again in the absence of procaine.

### Light-sensitivity test

Hym-176B::GCaMP transgenic animals without buds were first exposed to visible light for 1 min under a microscope to remove overly sensitive animals. The animals that did not contract in response to visible light were subsequently exposed to the blue excitation light, with or without a black filter to attenuate the intensity by 75%. We measured the time when the light-sensitive cell population started to glow under a microscope with 30x magnification. Since it usually took more than 2 min for the filtered blue light to trigger excitation, we removed the filter after 1 min exposure. Thus, for the filtered group, if the exposure time before excitation was more than 1 min, the filter was removed after 1 min and the population was thus exposed without the filter. Three independent experiments were conducted (n = 22 for exp_1 and 2, and n = 14 for exp_3). Statistical significance of the difference between the average time to the beginning of excitation with and without a dark filter was evaluated using twosided Student’s t-test with the R package, ggsignif.

---

**Movie 1. Hym-176A expressing neuron subset.** 4 to 8 sec of Movie S1 with higher magnification. Speed: 1 x real-time.

**Movie 2. Hym-176B expressing neuron subset.** 10 to 14 sec of Movie S2 with higher magnification. Speed: 1 x real-time.

**Movie 3. Hym-176C expressing neuron subset.** Full length of Movie S3 with higher magnification. Speed: 1 x real-time.

**Movie 4. Hym-176D expressing neuron subset.** Full length of Movie S4 with higher magnification. Speed: 1 x real-time.

**Movie 5. Double transgenic line; Hym-176B::GCaMP x Hym-176C::GCaMP.** Speed: 1 x realtime.

**Movie 6. Operative chimera; Hym-176B::GCaMP + Hym-176A::GCaMP.** Speed: 1 x real-time.

**Movie 7. Double transgenic line; Hym-176B::GCaMP x Hym-176C::GCaMP treated with procaine.** Speed: 1 x real-time.

**Movie 8. Operative chimera; Hym-176B::GCaMP + Hym-176A::GCaMP treated with procaine.** Speed: 1 x real-time.

**Movie 9. Hym-176B::GCaMP transgenic animal treated with procaine.** Speed: 1 x real-time.

**Movie 10. Hym-176B::GCaMP transgenic animal.** Speed: 4 x real-time.

## Notes

https://drive.google.com/drive/folders/1GngAsZTvRy4Kpba1Rq_YI9rPGQ_e7MEK?usp=sharing

### Competing Interest Statement

The authors have declared no competing interest.

