## Supplemental Data for "A single neuron subset governs a single coactive neuron circuit in *Hydra vulgaris*, representing a prototypic feature of neural evolution"

Fig. S1

| Gene# | Group | Gene | Peptide | Expression: Tentacle | Hypostome | Body column: upper | Body column: lower | Peduncle: upper | Peduncle: lower: upper | Peduncle: lower: lower | Basal disk |
|---|---|---|---|---|---|---|---|---|---|---|---|
| 1 | GLWa | GLWa | I, II, III, IV | v | v | v | v | v | v | v | v |
| 2 | Hym-355 | Hym-355 | Hym-355 | v | v | v | v | v | v | v | v |
| 3 | RFa | Prepro A | I, II, III, IV | v | v | v | | | v | v | |
| 4 | | Prepro B | I, II | | v | v | | | | | |
| 5 | | Prepro C | I | v | | | | | | | |
| 6 | Hym-176 | Hym-176A | Hym-176, Hym-357 | | v | v | v | v | v | v | |
| 7 | | Hym-176B | Hym-357 | | v | v | v | v | | | |
| 8 | | Hym-176C | (Hym-176, Hym-357) | | | | | | v | v | |
| 9 | | Hym-176D | (Hym-176) | | | | | | v | (v)[*3] | |
| 10 | | Hym-176E | – | v | | | | | | | |

| Classical subsets | Gene# |
|---|---|
| I | 1+2[*1] |
| II | 3+5 |
| III | 3+4 |
| IV | 6+7[*2] |
| V | 3+6+8+9 |
| VI | 3+6+8 |
| VII | 10 |

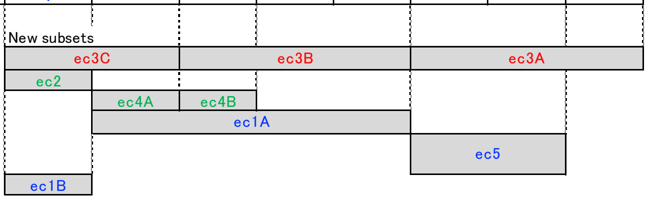


**Fig. S1. Mutually exclusive peptidergic neuron subsets in the ectodermal layer.** Upper panel: Neuropeptide genes distinguished by Gene# are expressed in neurons located in different regions of *Hydra*. Different gene expression shown in the same color in a given region indicates co-expression in the same neurons. Peptides in parenthesis are similar but not identical. Lower panel: According to the upper panel, mutually exclusive neuron subsets, each of which express a different combination of neuropeptide genes, are listed. The classical subsets express genes shown in the column, "Gene#". For example, neuron subset V expresses genes #3, #6, #8, and #9 in the lower peduncle. A recent single-cell RNA-seq study showed that the classical subset can be further divided into new subsets. *[1)] There are some GLWa$^{+}$/Hym355$^{-}$ neurons in the hypostome. [*2)] Expression of *Hym-176A* is much less than that of *Hym-176B* in the body column and the upper peduncle. [*3)] *Hym-176D* is not expressed in this region of strain 105 but expressed in strain AEP.

Fig. S2

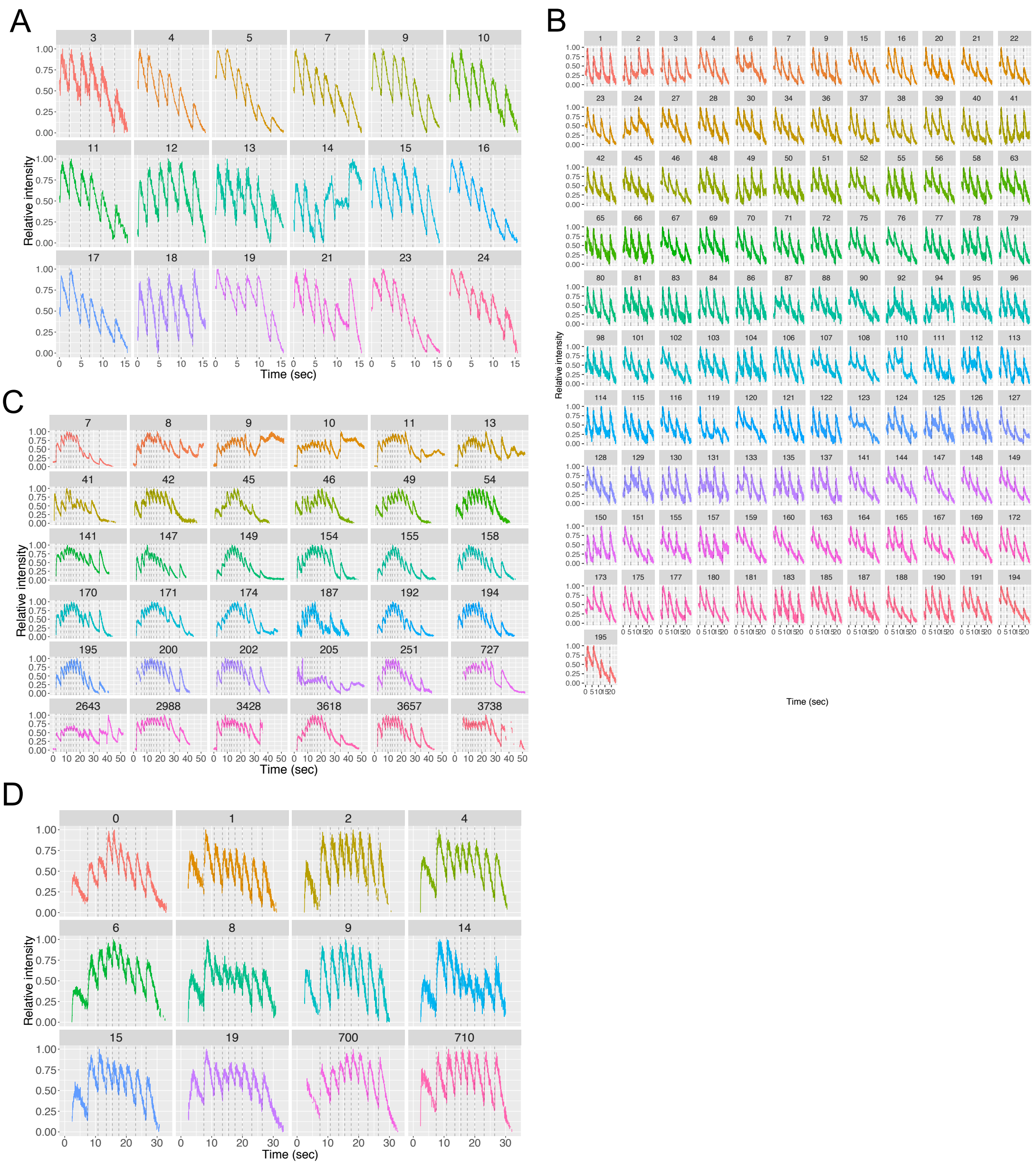


**Fig. S2. Normalized neuronal activity in the Hym-176 peptidergic neuron subsets.** *Hym-176A*-expressing neuron subset (A). *Hym-176B*-expressing neuron subset (B). *Hym-176C*-expressing neuron subset (C). *Hym-176D*-expressing neuron subset (D). Neuronal activity (normalized relative intensity of GCaMP) with vertical dashed lines indicate the average starting time of excitation of all tested neurons in each subset, as described in Fig. 1. The number in each strip and the color of the excitation profile are also described in Fig. 1.

Fig. S3

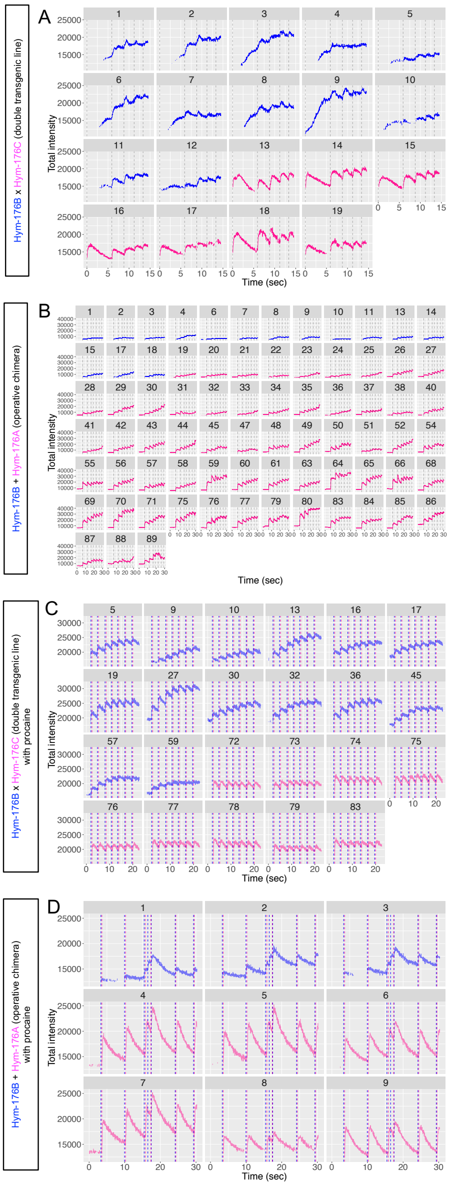

**Fig. S3. Unnormalized neuronal activity of the body and the foot neuron circuit.** Double transgenic line of Hym-176B::GCaMP and Hym-176C::GCaMP with (C) or without (A) procaine treatment. The operative chimera of Hym-176B::GCaMP and Hym-176A::GCaMP with (D) or without (B) procaine treatment. Unnormalized neuronal activity (total intensity of GCaMP) are plotted as described in Fig. 3B, D, F, H.

**Movie S1. Hym-176A-expressing neuron subset.** Speed: 4 x real-time.

**Movie S2. Hym-176B-expressing neuron subset.** Speed: 4 x real-time.

**Movie S3. Hym-176C-expressing neuron subset.** Speed: 4 x real-time.

**Movie S4. Hym-176D-expressing neuron subset.** Speed: 4 x real-time.

**Movie S5. Hym-176A, B, or D-expressing neuron subset.** High magnification. Speed: 4 x real-time.
